# GROQ-seq Enables Cross-site Reproducibility for High-Throughput Measurement of Protein Function

**DOI:** 10.64898/2026.04.07.716961

**Authors:** Aviv Spinner, David Ross, Dana Cortade, Svetlana Ikonomova, Catherine Baranowski, Andi Dhroso, Amanda Reider Apel, Kristen Sheldon, Courtney Duquette, Douglas Densmore, Peter J Kelly, Erika DeBenedictis, Corey M. Hudson

## Abstract

High-throughput functional assays are increasingly used to generate large-scale protein function datasets for protein engineering and machine learning applications. However, the utility of such datasets depends on the reproducibility of the underlying measurements. **Here we report reproducible, quantitative measurements of protein sequence-to-function data at scale across two facilities.** We analyze GROQ-seq (**G**rowth-based **Q**uantitative **Seq**uencing) measurements of three bacterial transcription factors. Independent barcode measurements of the same sequence produce highly consistent functional estimates, demonstrating strong biological reproducibility (across all transcription factors the mean Root Mean Square Deviation [RMSD] ≈ 0.53 and mean Spearman ≈ 0.63). We also compared experiments performed at two facilities using a shared protocol, but with differing levels of automation and system integration. We observe strong agreement between measurements taken at the two sites (mean RMSD ≈ 0.41 and mean Spearman ≈ 0.730). Orthogonal tests further support this agreement: a classifier trained to distinguish data by site performs near random (AUC = 0.559), and top-ranking variants show strong statistical overlap between experiments. Together, these results demonstrate that GROQ-seq enables reproducible, scalable measurement of protein function suitable for large aggregated datasets.

## Introduction

Large-scale datasets are essential for building accurate and generalizable AI models in protein science^1^. However, generating data that are both sufficiently large and consistently high quality remains a major challenge. Detailed studies of individual proteins have enabled protein sequence–function mapping and the development of protein-specific models^2–5^. As datasets expand^6^ and model architectures become more transferable^7,8^, the field has shifted toward generalizable predictive models^9–13^. However, the field’s progression has also exposed limitations in data quality and consistency^14^. Unlike genomics^15–17^ and protein structure prediction^1,2,18,19^, which benefit from decades of standardized data accumulation, protein function datasets remain bespoke. Traditionally, each new protein function requires a custom assay, and few assays are reused across studies, resulting in small, fragmented datasets^20,21^. As a result, progress toward broadly applicable predictive models is limited by the difficulty of integrating measurements across experiments and studies.

Reproducible data generation would enable scalable dataset aggregation for machine learning. Reproducibility can be defined at several levels (Box 1), including technical, biological, protocol, and cross-site reproducibility. Assays must provide sufficient dynamic range and well-characterized error to distinguish true biological effects from noise. These challenges are particularly pronounced in pooled growth-based assays, where thousands of variants are measured simultaneously and small biases in growth, amplification, sampling, or sequencing can compound over time. Although some variability is inherent to biological systems, systematic bias is especially problematic, as it introduces consistent shifts between experiments that cannot be averaged out^22^. Consequently, both within-experiment biological reproducibility and cross-site reproducibility are essential for generating reliable and scalable datasets for machine learning.

Here, we analyze the reproducibility of GROQ-seq, a high-throughput pooled assay designed to quantitatively measure protein function at scale. We evaluate the reproducibility of GROQ-seq both within experiments and across independent facilities. We find that GROQ-seq produces highly reproducible measurements across experimental replicates and sites, demonstrating its potential as a robust platform for generating scalable, standardized, and AI-ready datasets for protein function prediction.

**Box 1:**
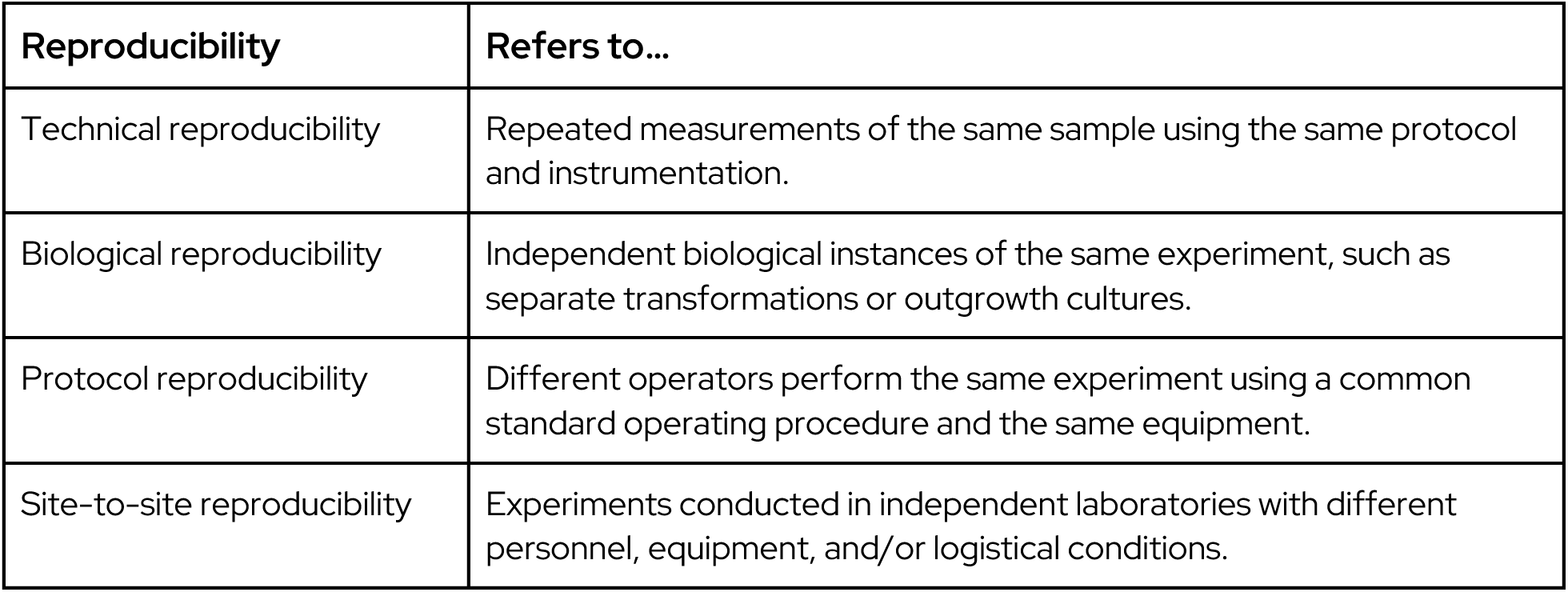
Definitions of reproducibility.

### GROQ-seq Assay

GROQ-seq is a pooled growth based assay that enables quantitative measurement of sequence-to-function datasets^23^. Briefly, GROQ-seq works by coupling the function of a protein to growth of a bacterial cell through the use of a genetic circuit or auxotrophy, enabling pooled measurement of barcoded protein variant fitness (Figure 5). The method has been used in the past to measure libraries of 100k protein variants in a single pooled experiment^24,25^. GROQ-seq differentiates itself through the use of an internal calibration ladder of variants with known functional values to enable two advances: improved consistency across batches and facilities, and conversion of enrichment measurements into quantitative, system-specific functional units (e.g., *k*_cat_). GROQ-seq circuits have been built for transcription factors, TEV Protease^26^, and T7 RNA Polymerase^27^ and have been proposed to be compatible with many additional protein function families^28^, making it a general-purpose approach to collection of large sequence to function datasets.

Here we analyze biological and site-to-site reproducibility results of GROQ-seq measurements of transcription factor function on three bacterial transcription factors (RamR, LacI, and VanR) in *Escherichia coli*. In GROQ-seq, the function measured depends on the protein class; for transcription factors, this corresponds to regulation of gene expression (transcription). The transcription factor binds to its cognate DNA operator and thereby regulates the transcription of a dihydrofolate reductase (DHFR) gene (Supplemental Figure 1). DHFR modulates bacterial resistance to trimethoprim (TMP) and thus growth^29,30^. To quantify the response of each TF to its small-molecule ligand, we use the GROQ-seq assay to measure two function values: the uninduced transcription rate (without ligand) and the induced transcription rate (with a high concentration of ligand). In general, the transcription rate varies over approximately 2.5 orders of magnitude, and we chose the calibration ladder variants to span that full range. To assess reproducibility, we analyze GROQ-seq data for the two calibrated function values and their ratio (i.e., the induced to uninduced transcription rate ratio). We analyze data including measurements of site saturation variant libraries (SSVL), site saturation mutagenesis (SSM), and error-prone PCR (epPCR) libraries of transcription factors (Methods).

### Biological Reproducibility

We leverage barcode redundancy to evaluate biological reproducibility within each GROQ-seq experiment. A central challenge in high-throughput biological measurement is ensuring that observed functional differences reflect true biological effects rather than stochastic experimental variation. In pooled growth assays like GROQ-seq, hundreds of thousands of variants may be assayed simultaneously and this challenge is particularly acute. Individual amino acid variants associated with multiple different DNA barcodes allow for the same amino acid sequence to be observed through multiple independent measurements within a single experiment. Because each barcode represents an independent transformation and lineage within the pooled population, agreement between barcodes provides a direct estimate of biological variation arising during growth and selection.

Across the RamR library, 18.85% of amino acid variants are represented by two or more independent barcodes (Figure 1A), and we can assess the biological variability by comparing their function measurements. The distribution of barcode-to-barcode variability across sequences is shown in Figure 1B. The median standard deviation across barcodes is approximately 0.2 (in base-10 log units), corresponding to ≅1.6-fold variation in functional measurements. This level of variability is small compared to the 2.5-fold dynamic range of the assay, indicating low measurement noise.

**Figure 1:**
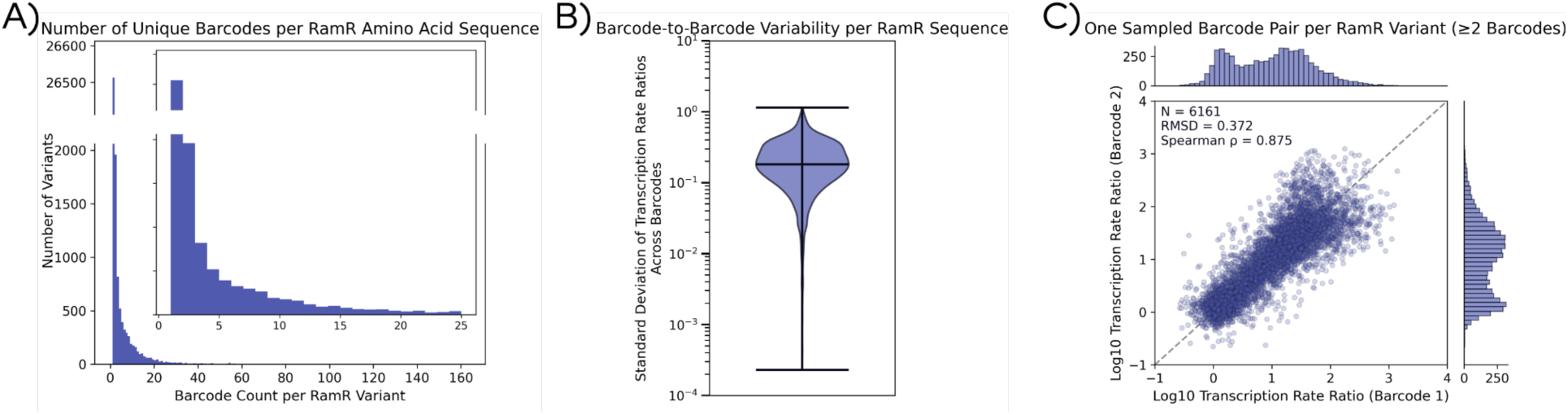
Biological reproducibility within a GROQ-seq experiment. **(A)** Distribution of barcode counts per amino acid variant in the RamR library. The insert plot focuses on up to 25 barcode counts. The majority of protein variants correspond to a single barcode, while 18.85% of variants are associated with multiple independent barcodes that serve as internal biological replicates. **(B)** Biological variability for amino acid variants with multiple barcodes was quantified as the standard deviation of the log10 transcription rate ratio (ratio of the induced to uninduced transcription rate) among barcodes associated with the same variant. The median standard deviation of ≅0.2 corresponds to 10^0.2^ ≅1.6-fold variation. **(C)** Agreement between independent barcode measurements for the same variant. Each point represents a randomly sampled pair of barcodes corresponding to the same amino acid sequence (N = 6,161 pairs), demonstrating strong concordance across these measurements (RMSD = 0.372, Spearman = 0.875). The marginal distributions are shown on the top and right axes.

Next, we assessed whether independent barcode measurements for the same sequence produce consistent functional estimates across the measurement’s dynamic range. Barcode-level differences could arise from several factors, including differences in transformation efficiency, differential growth dynamics within the pooled culture, PCR amplification bias during library preparation, or variation in sequencing depth. For variants represented by at least two barcodes, we randomly sampled pairs of barcodes and compared the resulting measurements (Figure 1C). To appropriately assess reproducibility, we report both Root Mean Squared Deviation (RMSD) and Spearman correlation (Methods: Statistic). These measurements show strong agreement, with an RMSD of 0.372 (2.4-fold) and a Spearman correlation of 0.875 across 6,161 sampled barcode pairs. The strong correlation between sequences tagged with independent barcodes demonstrates that functional measurements are largely determined by the underlying amino acid sequence rather than stochasticity in measurement.

Identical analysis is conducted for two additional transcription factors, LacI and VanR as well as all three transcription factors tested at DAMP (Supplement Figures 2-6). Across all proteins and experimental sites, we observe similarly strong agreement between independent barcode measurements, with Spearman correlations consistently in the range of ∼0.6–0.8 and RMSD values indicating low measurement variability, demonstrating that these results generalize across diverse proteins and experimental conditions and that GROQ-seq measurements are highly reproducible within a single pooled experiment. This level of biological reproducibility provides a critical foundation for interpreting downstream analyses, including comparisons across experiments and between independent laboratories.

### Site-to-site reproducibility

Next, we analyzed reproducibility across independent laboratories, which represents a substantially stronger test of assay robustness. Differences in personnel, instrumentation, automation workflows, protocol reagents, and sequencing depth can introduce unintentional bias and noise in pooled growth assays. Establishing that GROQ-seq measurements remain stable across such differences is critical for enabling comparisons across institutions and for supporting long-term data scaling and aggregation of datasets.

To assess site-to-site reproducibility, the GROQ-seq assay was independently performed at two facilities: the Living Measurements System Foundry (LMSF) at the National Institute of Standards and Technology (NIST) in Gaithersburg, Maryland; and the Design, Automation, Manufacturing, and Processes (DAMP) laboratory at Boston University in Boston, Massachusetts. Although both experiments followed the same standard operating procedure, several operational differences existed between sites (Table 1). The experiments were initiated from separate outgrowths of a shared glycerol stock, meaning that the initial composition of the pooled population was expected to be similar, but not identical between the two sites.

**Table 1:** Site-to-site laboratory comparison between DAMP and LMSF.

| GROQ-seq Step | DAMP | LMSF |
| --- | --- | --- |
| Overnight inoculation | manual, benchtop |  |
| Plate setup | Hamilton STAR |  |
| Plate incubation (pre-warming) |  |  |
| Plate inoculation |  |  |
| Plate movement | Manual | Integrated Robotic Arm |
| Plate sealing/de-sealing | Manual | Azenta a4S Heat Sealer and<br>Azenta XPeel desealer |
| Plate Incubation | BioTek Synergy H1 plate reader<br>37°C, double-orbital shaking at 807<br>cycles per minute | BioTek Synergy Neo2 plate reader<br>37°C, double-orbital shaking at<br>807 cycles per minute |
| Plasmid DNA extraction | Hamilton STAR - bead based | Hamilton STAR - column based |
| BarSeq Liquid handling | Hamilton STAR |  |
| BarSeq thermocycler | Off-deck | On-deck |
| BarSeq cleanup | Hamilton STAR - bead based |  |
| BarSeq cleanup method |  |  |
| BarSeq QC | Tapestation, QuBit | Gels, QuantIT |
| Lab setup | Open environment | Integrated single PAA S-Cel<br>Workstation |

Sequencing depths also differed between experiments: The LMSF samples were sequenced with a 10B-read Illumina NovaSeq flow cell followed by a second, 25B-read flow cell to improve depth of coverage for low-abundance barcodes, yielding a total of 19.9B reads. The DAMP samples were sequenced on a single 25B-read flow cell yielding 4.5B reads. Additionally, the DAMP lab uses experimental equipment in an open environment while the LMSF equipment is contained within an integrated PAA S-Cel Workstation. These differences create potential sources of variation in measured barcode counts and variant representation across sites.

Despite these sources of variation, the two experiments assayed a large shared set of amino acid variants. Due to differences in sequencing depth, the total number of reported variants, defined as variants passing minimum read count thresholds during initial data filtering, was 32,592 at LMSF and 16,159 at DAMP. Of the variants observed across the two datasets, 15,383 were present in both experiments (Figure 2A). The distribution of barcode counts per shared variant reflects differences in sequencing depth and sampling between the sites, with LMSF generally exhibiting higher barcode counts due to deeper sequencing (Figure 2B).

**Figure 2:**
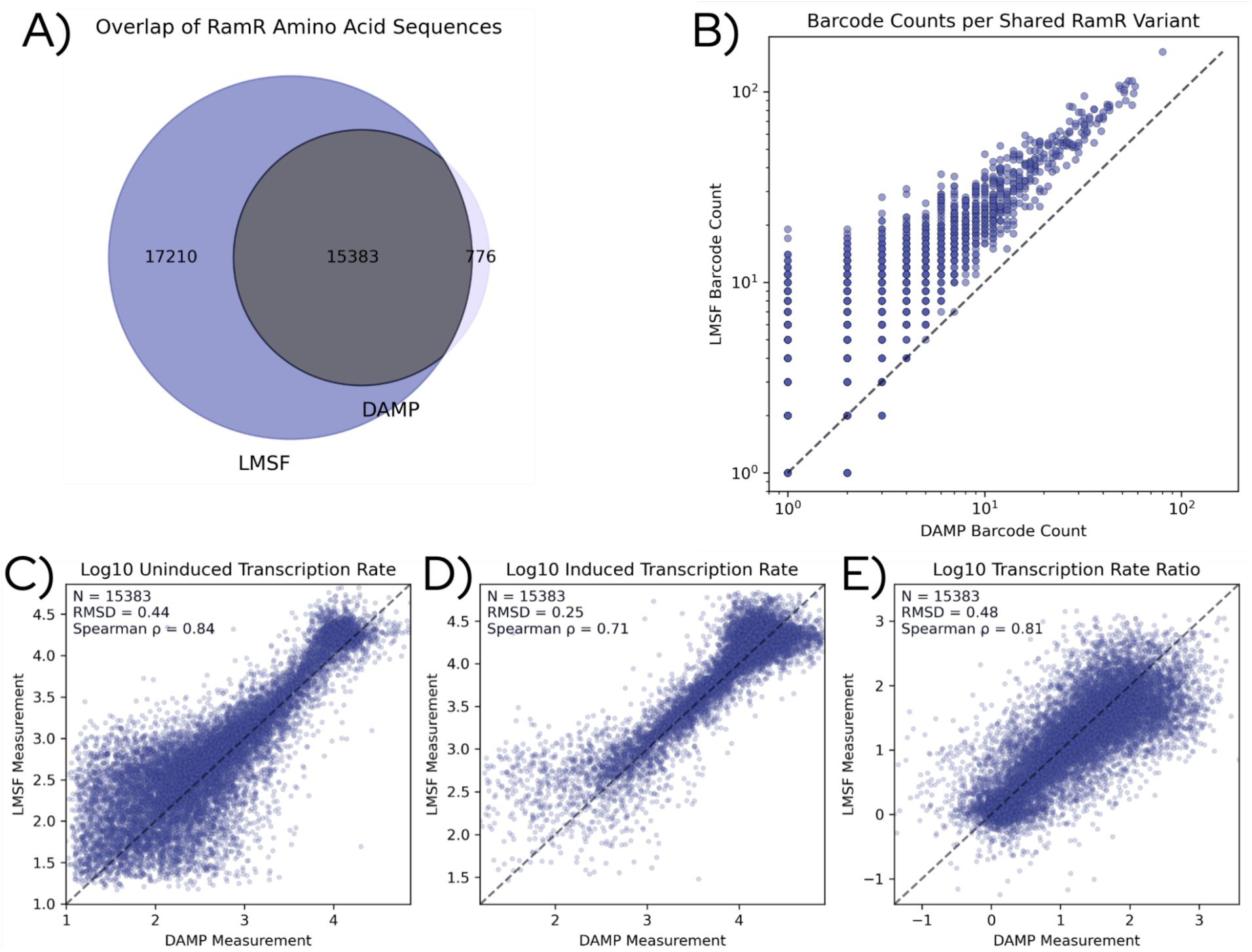
Site-to-site reproducibility of GROQ-seq measurements between LMSF and DAMP laboratories. **(A)** Overlap of amino acid variants measured at each site for the RamR library. Of all observed variants, 15,383 were shared between datasets, with additional variants detected uniquely at LMSF (17,210) or DAMP (776). **(B)** Comparison of barcode counts for shared variants across sites, showing broadly consistent variant representation despite differences in sequencing depth. **(C–E)** Comparison of functional measurements for variants observed at both sites. Scatter plots show agreement between DAMP and LMSF measurements for the log10 uninduced transcription rate **(C)**, log10 induced transcription rate **(D)**, and log10 induced-to-uninduced transcription rate ratio of calibrated functional measurements **(E)**. Uncalibrated measurement analysis can be found in Supplemental Figure 7.

To evaluate reproducibility across laboratories, we compared the functional measurements for variants shared between the two sites. We compared three functional measurements derived from the assay: the uninduced transcription rate, the induced transcription rate, and the ratio between these two measurements, which captures the functional response of each variant to induction. Across all three measurements, we observe strong agreement between the two sites (Figure 2C–E). Similar trends are observed for uncalibrated fitness measurements, which show comparable rank-order agreement across sites (Supplementary Figure 7). The uninduced condition exhibits an RMSD of 0.44 with a Spearman correlation of 0.84, while the induced condition shows an RMSD of 0.25 with a Spearman correlation of 0.71. Because these RMSD values are calculated on log10-transformed measurements, they correspond to differences of approximately 2.75-fold RMSD and 1.78-fold RMSD between experiments, respectively. The induced-to-uninduced ratio, which represents an important performance parameter when transcription factors are used as biosensors, exhibits an RMSD of 0.48 and a Spearman correlation of 0.81, corresponding to a typical difference of approximately 3.0-fold RMSD between measurements across sites. Variants maintain highly consistent functional measurements across independent laboratories despite differences in sequencing depth and experimental details.

Beyond pairwise comparisons of individual variants, we also examined whether the overall distribution of functional measurements was consistent between sites. The distributions of functional scores from the two experiments show nearly identical shapes, indicating that the global structure of the protein function landscape is preserved across laboratories (Figure 3A). This agreement is notable, as differences in variant representation between sites, along with potential biases from PCR amplification, growth dynamics, and/or sequencing depth, could all distort the calculated global structure of the protein functional landscape. The close match between distributions suggests that these effects are minimal and that both experiments recover the same underlying biological signal rather than site-specific artifacts. We trained a logistic regression classifier using functional measurements labeled by their site of origin, using a balanced dataset with equal representation from each site. If systematic differences existed between the two experiments, such a classifier should be able to distinguish measurements from LMSF and DAMP. However, when trained using an 80/20 train–test split, the classifier performs only marginally better than random guessing, with an area under the receiver operating characteristic curve (AUC) of 0.559 (Figure 3B). The inability of the classifier to distinguish between measurements from the two laboratories further supports the conclusion that the experiments produce nearly indistinguishable functional landscapes.

**Figure 3:**
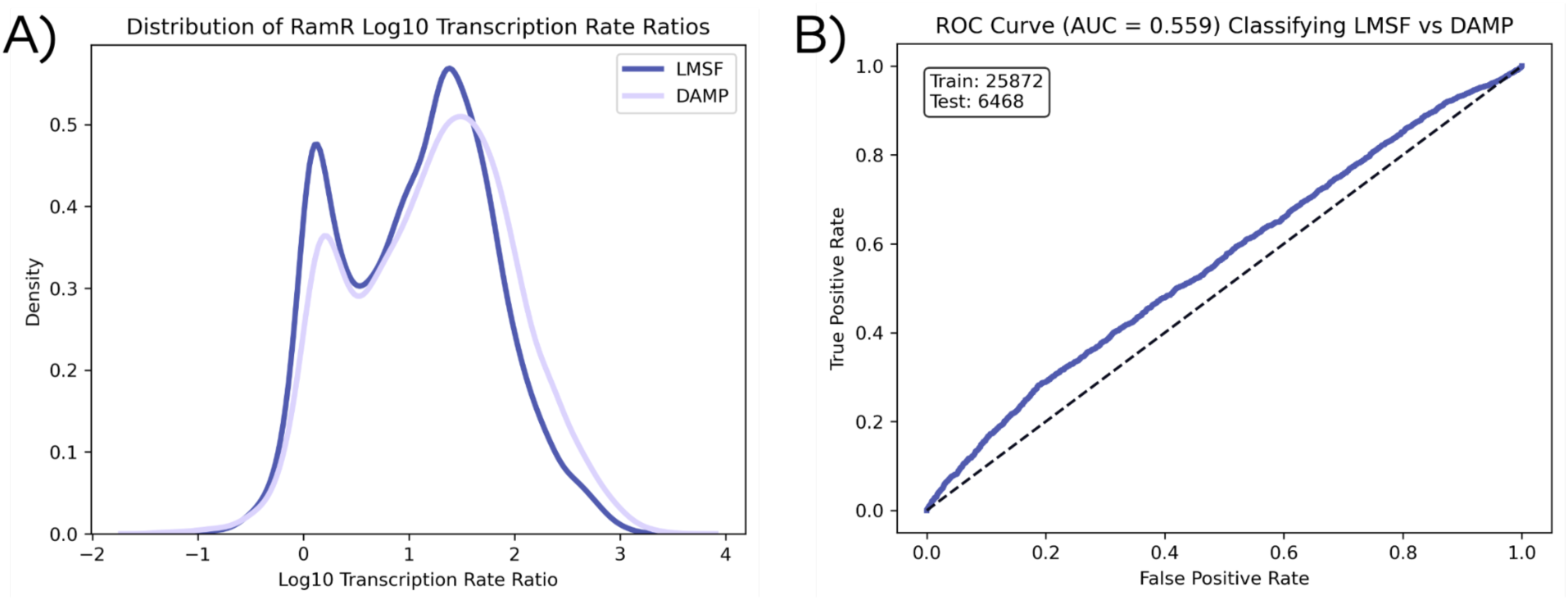
Global similarity of functional landscapes across experimental sites. **(A)** Distribution of Log10 transcriptional rate ratio measurements from LMSF and DAMP, showing highly similar overall distributions of variant effects across the two experiments. The differences in height between these distributions is likely due to differences in sequencing depth rather than underlying biological differences. **(B)** Logistic regression classifier trained to distinguish variants measured at LMSF versus DAMP performs near random (AUC = 0.559) with a train/test split of 80/20 using log10 transcriptional rate ratio measurements. Results here indicate that the global structure of the functional landscape is indistinguishable between sites.

Finally, we examined reproducibility at the extreme end of the functional distribution. For many applications in protein engineering and biological discovery, accurately identifying variants at the tails of the distribution can be more important than achieving perfect agreement across the entire population. For transcription factors engineered as biosensors, variants with larger log10 transcriptional rate ratios are generally preferred because they produce stronger and more distinguishable responses between induced and uninduced conditions. Under this definition, variants appearing at the upper end of the functional distribution correspond to those with the largest functional responses under this metric. If the assay is reproducible, variants with the largest functional response at one site should also have the largest functional response when measured independently at another site.

To evaluate this, the overlap between the top-ranked variants from the LMSF and DAMP experiments was compared. For each value of *N*, the number of variants shared between the top *N* variants from each dataset (Figure 4A) was computed. For example, for *N* = 100, we asked how many variants are shared between the top 100 variants in LMSF and DAMP. Then, the observed overlap was compared to the overlap expected under random sampling using a hypergeometric distribution. If *M* is the total number of shared variants between the two datasets, and each site contributes a set of size *N*, then the number of shared variants *X* under random sampling follows:*X ∼ Hypergeometric(M, N, N)*

**Figure 4:**
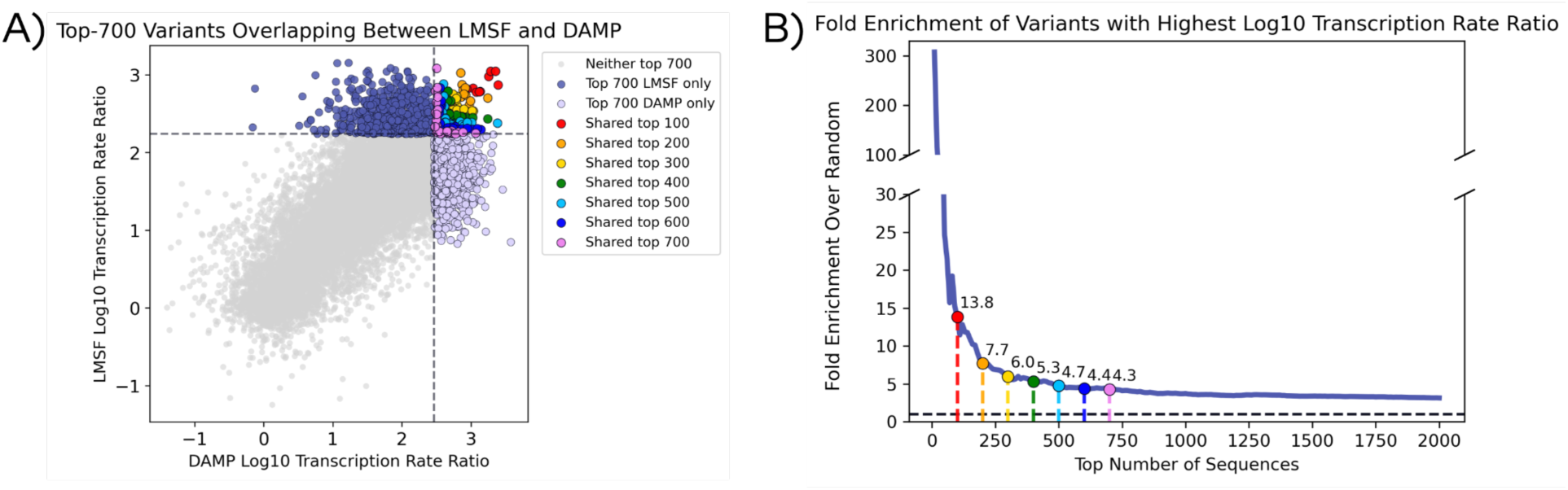
Reproducible identification of highly functional variants across experimental sites. **(A)** Scatter plot of Log10 transcriptional rate ratio for variants shared between LMSF and DAMP experiments, highlighting variants that fall within the top-N highest functional scores at both sites. Colored points indicate variants shared among the top-ranked sets (top 100–700), demonstrating strong agreement in the identification of highly and moderately functional variants. **(B)** Fold enrichment of shared top-N variants between LMSF and DAMP relative to the overlap expected under random sampling of the hypergeometric distribution. The observed overlap is strongly enriched across a wide range of N, indicating that variants identified near the extreme end of the functional distribution are reproducibly recovered across laboratories.

The expected overlap under this null model is 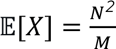. Fold enrichment is the ratio of the observed overlap to the expected overlap under random sampling where 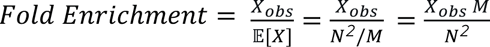. For example, considering the top 20 variants from each dataset, *N* = *20*, and *M* = *15*,*383* (total number of variants), under a random-sampling assumption, the expected number of shared variants between the two top-20 lists is: 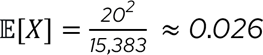. So, with random sampling, it is expected that essentially zero shared variants exist between the two top-20 lists. In data presented here, there is an observed overlap of *X_obs_*, = *3* therefore 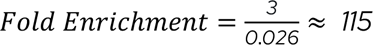. For the list of top 100 variants, the fold enrichment is almost 14-fold. Across a wide range of values of *N*, the overlap between top variants remains several-fold higher than expected by chance even for large values of *N* (Figure 5B). Therefore, GROQ-seq reliably identifies highly functional variants and, more broadly, recovers moderately functional variants at rates above random expectation.

**Figure 5:**
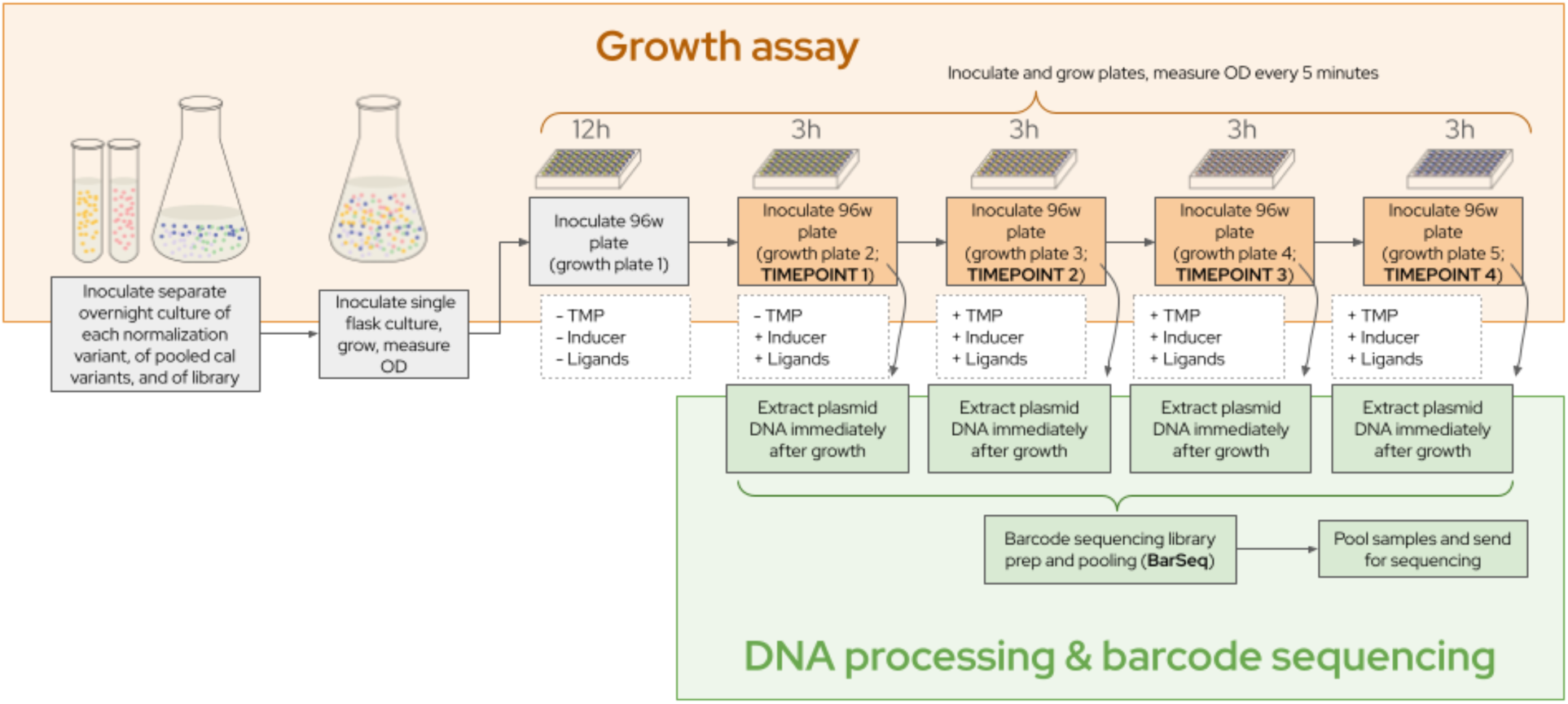
Schematic overview of the GROQ-seq growth assay and sequencing workflow. Barcoded protein variant libraries are grown in pooled culture across timepoints under defined induction and selection conditions, with plasmid DNA extracted after each growth phase to capture changes in variant abundance. These measurements are converted into quantitative functional readouts through barcode sequencing (BarSeq) and internal calibration, enabling high-throughput, mapping of sequence–function relationships.

Identical analysis was performed for the other two transcription factors, LacI and VanR (Supplemental Figures 7-13). Across both additional proteins, there is similarly strong agreement between sites, with consistent rank-order concordance (Spearman ∼0.5–0.9), modest RMSD values, near-random AUCs, and significant fold enrichment of top variants, indicating that these results generalize across proteins. Agreement between laboratories is far greater than would be expected by chance alone, providing strong statistical evidence that GROQ-seq faithfully reproduces the highest-performing variants across independent experimental sites.

## Methods

Please refer to the “Technical Bulletin for GROQ-seq Transcription Factor Function Assay” for an in-depth description of methods^30^.

### Variant Library Sourcing

Three types of variant libraries were pooled together and experimentally characterized using GROQ-seq for each transcription factor (RamR, VanR, and LacI): saturation variant libraries (SSVLs), site-saturation mutagenesis (SSM) libraries, and error-prone PCR (epPCR) libraries (Supplemental Figure 14). SSVLs were purchased from Twist Bioscience (South San Francisco, California) and included all single amino acid substitutions, deletions, and insertions across the full length of each protein. The SSM libraries were produced in-house at LMSF and included all combinatorial amino acid mutations focused on three residues involved in transcription factor–DNA interactions^25^. The epPCR libraries were purchased from GenScript USA Inc. (Piscataway, NJ) and contained variants with mutations randomly distributed across the protein sequence.

### Plasmid library construction and QC

Protein variant libraries and random barcodes were cloned into the GROQ-seq TF backbone plasmid (Supplemental Figure 1). For the SSVL and epPCR libraries this process was performed by Clutch Biotechnologies (Burlingame, California) using Golden Gate Assembly, while the cloning for the SSM libraries was performed in-house at LMSF using Gibson Assembly. The backbones (pTwist Kan Medium Copy from Twist Biosciences, South San Francisco, CA) contain a p15A origin of replication, a kanamycin resistance marker and a DHFR gene. Driving expression of the DHFR gene is a ligand inducible promoter specific to the TF of interest (pTac for LacI, pVanCC for VanR, and pRamR for RamR). This promoter contains within, or proximally, operator sites specific to the TF of interest. Barcodes were mapped to variants via long-read sequencing using Oxford Nanopore Technology (ONT) via Plasmidsaurus (Plasmidsaurus Inc., USA) and libraries were transformed into *E. coli* lacking the lac operon *(MG1655Δlac*^31^).

### Growth-assay

The growth assay measures cell abundance across four timepoints while cells are maintained in mid-log phase to ensure a near constant growth rate (Figure 5). To start, the TF variant libraries were grown overnight, alongside normalization and TF calibration strains. These were subcultured into a combined flask for one doubling, then used to inoculate the first plate of the GROQ-seq assay. Growth in the first plate is used to acclimate the cells to plate-based growth conditions and is not included in subsequent analysis. After 12 hours of growth, these cells were subcultured into growth plate (GP) 2, which contained the TF specific ligand [Isopropyl β-D-1-thiogalactopyranoside (IPTG) for LacI, (S)-1,2,3,4-Tetrahydroisoquinoline (1S-TIQ) for RamR and Vanillic Acid for VanR]). GP2 was grown for 3 hours before being used as inoculum for GP 3 (containing TMP and ligands) while the remaining cells from GP 2 were used for plasmid DNA extraction and served as the timepoint 1 measurement. This process—3 hours of growth followed by subculturing into the next plate and plasmid DNA extraction—was repeated for GPs 3–5, corresponding to timepoints 2–4. GPs 3–5 contained both TF-specific ligands and TMP, thereby applying the selective pressure required to assay protein variant function.

### Calibration

The calibration ladder for this dataset is composed of 8 LacI, 10 RamR and 4 VanR variant/operator combinations with both higher and lower basal (G0) and induced output compared to wild-type. The calibration variants were chosen to evenly span the full dynamic range of the assay. The TF pilot dataset shows approximately 2.5 orders of magnitude dynamic range (<10^2^ to >10⁴), defined as the region where differences in function can be quantitatively measured with GROQ-seq. A large dynamic range is important for capturing the range of regulatory activities observed in most functional screens and reliably measuring quantitative differences across that range. We also want to avoid saturation of highly active sequences, preserve resolution among mid-range sequences, and distinguish weak, moderate, and strong effects. Finally, an appropriate dynamic range allows for reliable normalization, comparison, and statistics across conditions.

### Barcode sequencing (BarSeq)

Plasmid libraries were extracted from all wells of GP 2-5 using either a column or bead based method. Extracted DNA was used as the template for BarSeq, a PCR protocol for preparing barcodes from the pooled growth assay plasmid DNA for sequencing (Novogene Co., Ltd using the Illumina NovaSeq X platform, paired-end 150 bp) to quantify differential variant fitness as growth rates. Please see the TF Technical Bulletin for BarSeq protocol^30^.

### Statistic

We use the root mean square deviation (RMSD) as the primary metric to assess reproducibility, as it directly quantifies the magnitude of differences between measurements across experiments. Correlation coefficients (e.g., Spearman, Pearson) are commonly reported as measures of agreement because they capture the degree to which variants maintain consistent rank ordering. However, correlation is not a direct measure of reproducibility, as it depends strongly on the underlying distribution of the data and can conflate measurement consistency with dynamic range. For example, if all variants have nearly identical measurements, even highly reproducible experiments can yield low correlation values. For comparability with prior protein sequence–function studies, we also report Spearman correlation.

### Data Analysis

Barcode sequencing data were analyzed following workflows similar to those described for previous work^24^. Briefly, the barcode read counts were normalized by the read count for high-fitness “normalization control” plasmid (with constitutive DHFR expression) to correct for differences in DNA extraction and PCR efficiency across different samples. The change in the normalized read count over the four time points was used to determine the fitness associated with each barcoded plasmid. The fitness values for 22 calibration ladder plasmids were used with a flow-cytometry-based calibration datasheet to create a calibration curve for the GROQ-seq dataset. The calibration curve was used to estimate the function for every barcoded TF variant via Bayesian inference with Markov chain Monte Carlo (MCMC) using the cmdstanpy interface to Stan^32^(https://mc-stan.org/docs/reference-manual/).

### Conclusion

High-throughput functional assays have transformed protein engineering and biological discovery. However, the value of such datasets depends critically on their reproducibility. In this work, we demonstrate that GROQ-seq produces highly consistent functional measurements both within experiments and across independent laboratories. Independent barcode measurements of the same variant yield well-calibrated and near-identical results, and experiments performed at separate sites recover nearly indistinguishable functional landscapes and identify the same highly functional variants.

Beyond validating a single assay, these results highlight the importance of designing high-throughput experiments with reproducibility and quantitative dynamic range as central goals. Reliable measurements with large dynamic range allow meaningful distinctions between weak, moderate, and strong functional effects while minimizing systematic bias across experiments.

Importantly, the quality and variance structure of experimental data can strongly influence the performance of machine learning models trained on biological measurements. Recent work in reinforcement learning has demonstrated that the statistical properties of training data, including variance and dynamic range, can significantly affect model performance and generalization. By producing reproducible measurements with well-calibrated variance and large dynamic range, assays such as GROQ-seq provide a strong foundation for building large, reliable datasets that can power the next generation of machine learning models for protein function.

## Acknowledgements

This work was supported by Griffin Catalyst and Schmidt Sciences. The DAMP Lab acknowledges laboratory support from its industrial members including Hamilton, OpenTrons, MITRE, SciSure, New England BioLabs, Transfyr, and SiLA.

## Disclaimer

Certain commercial equipment, instruments, or materials are identified to adequately specify experimental procedures. Such identification implies neither recommendation or endorsement by the National Institute of Standards and Technology nor that the materials or equipment identified are necessarily the best available for the purpose.

## Supplemental Figures

**Supplemental Figure 1:**
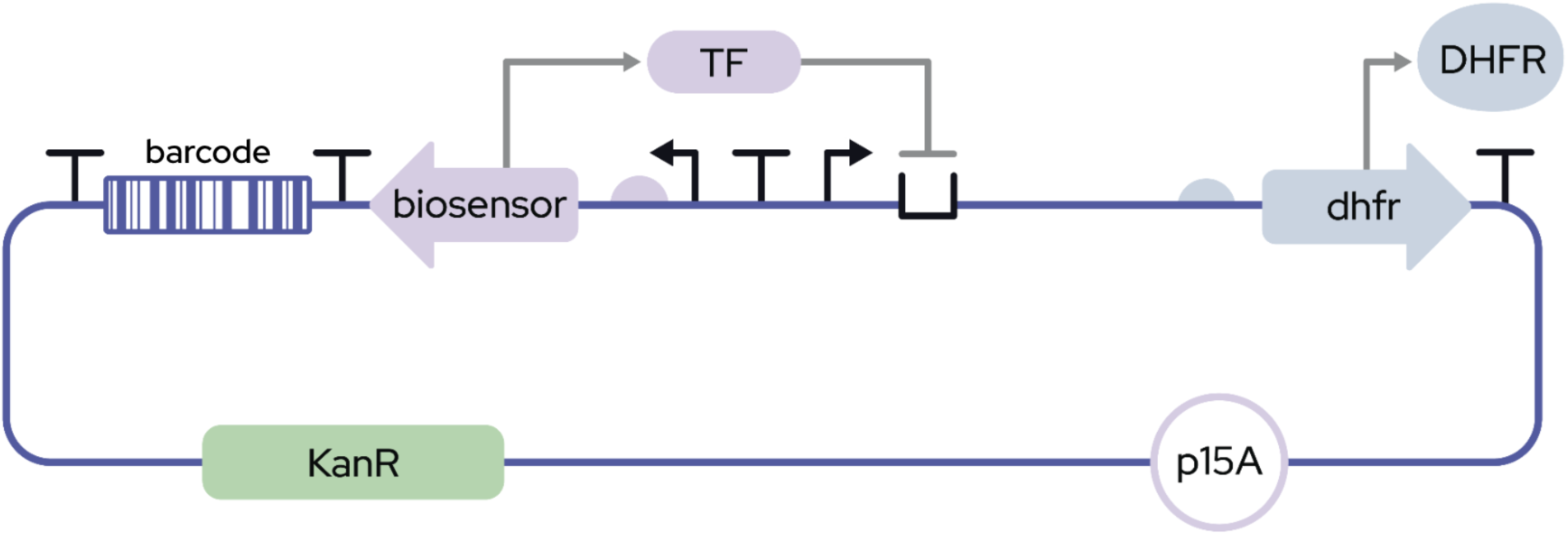
Circuit design used for GROQ-seq transcription factor measurements^29^. The gene of interest encodes a transcription factor (TF) that represses expression of a downstream DHFR reporter; stronger binding between the TF and the DNA operator results in decreased DHFR expression, which decreases resistance to trimethoprim and impacts cellular growth.

**Supplemental Figure 2:**
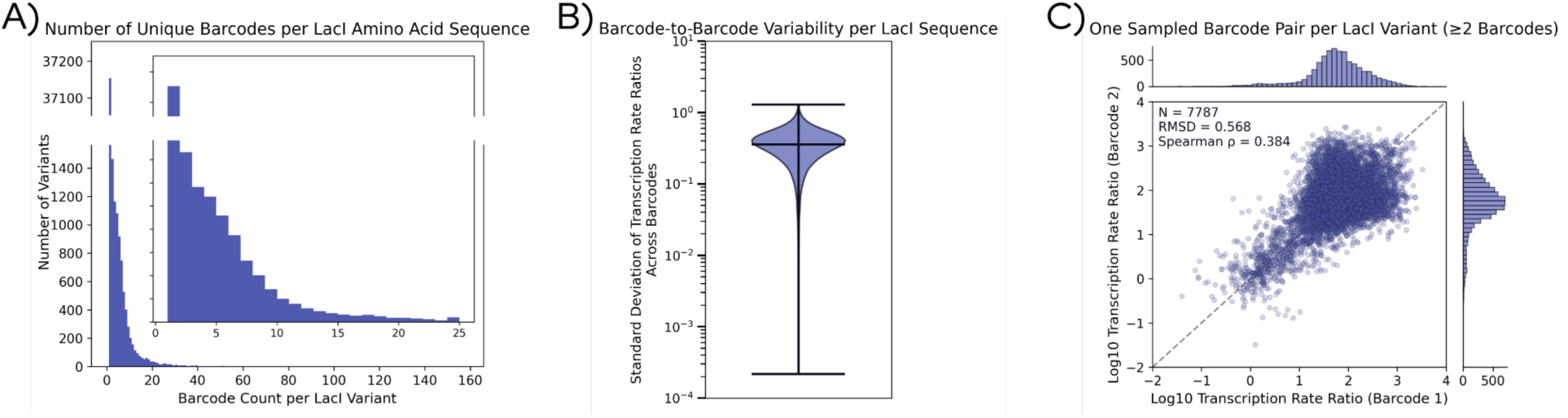
Corresponding main text Figure 1 for LacI at LMSF.

**Supplemental Figure 3:**
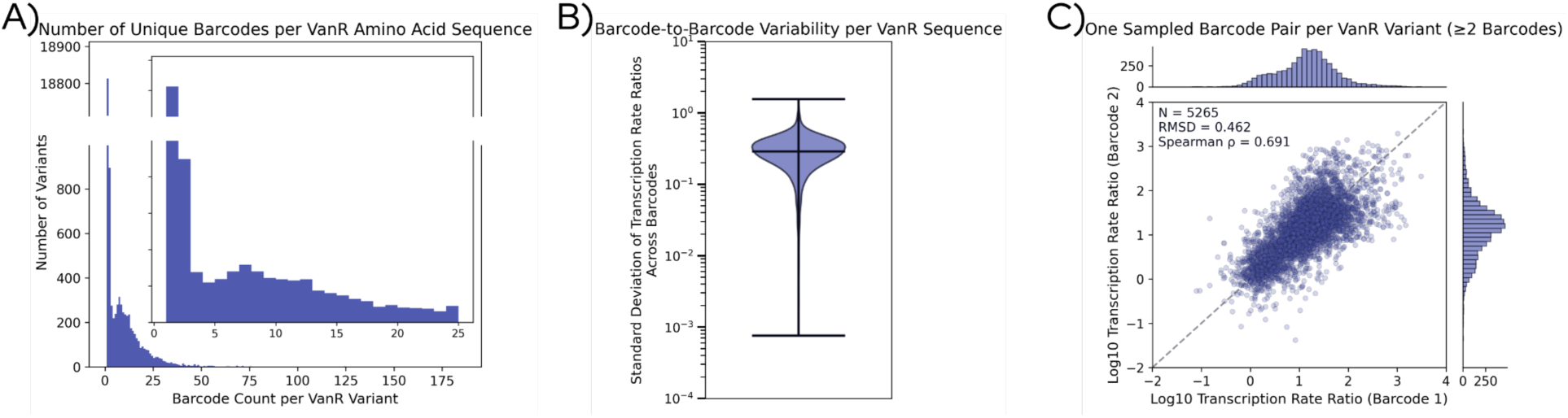
Corresponding main text Figure 1 for VanR at LMSF.

**Supplemental Figure 4:**
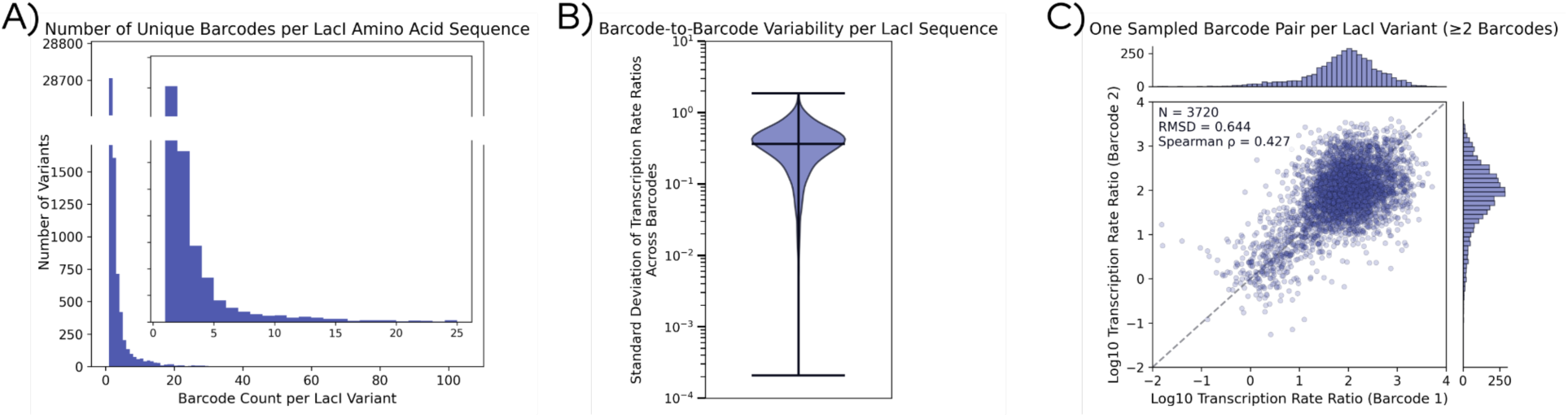
Corresponding main text Figure 1 for LacI at DAMP.

**Supplemental Figure 5:**
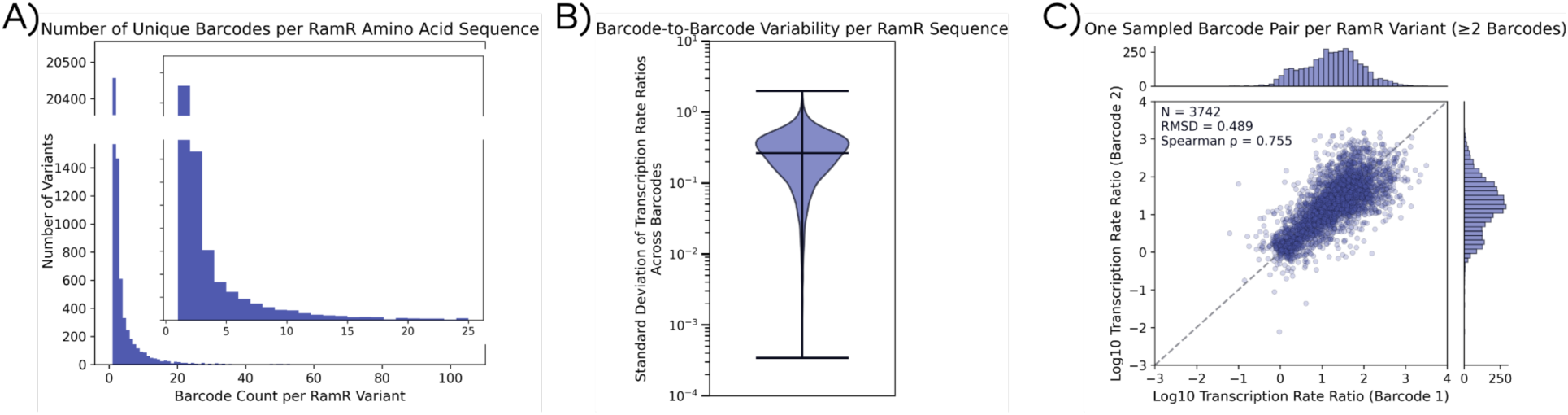
Corresponding main text Figure 1 for RamR at DAMP.

**Supplemental Figure 6:**
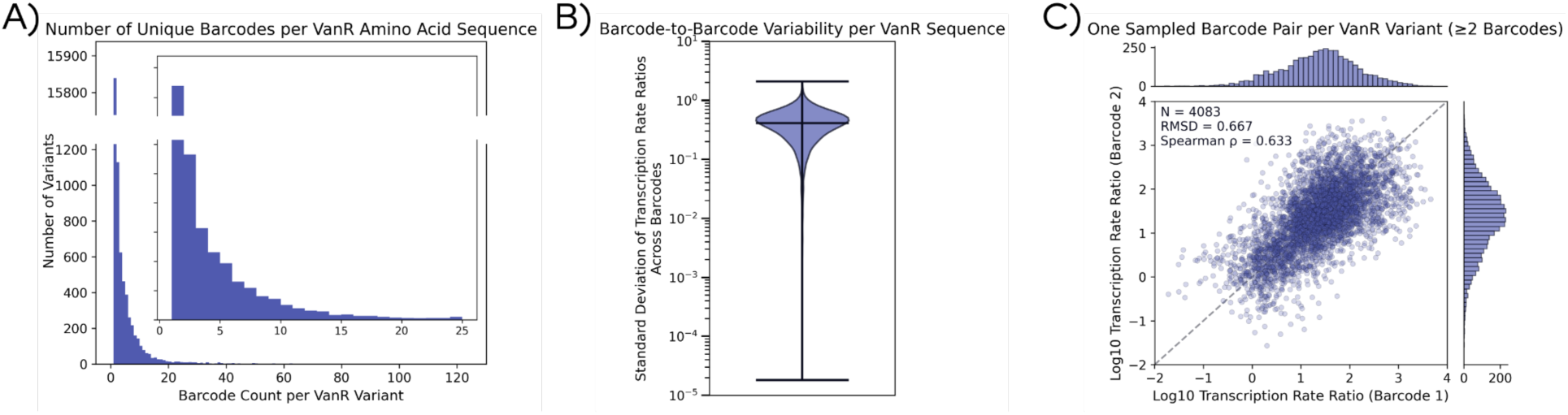
Corresponding main text Figure 1 for VanR at DAMP.

**Supplemental Figure 7:**
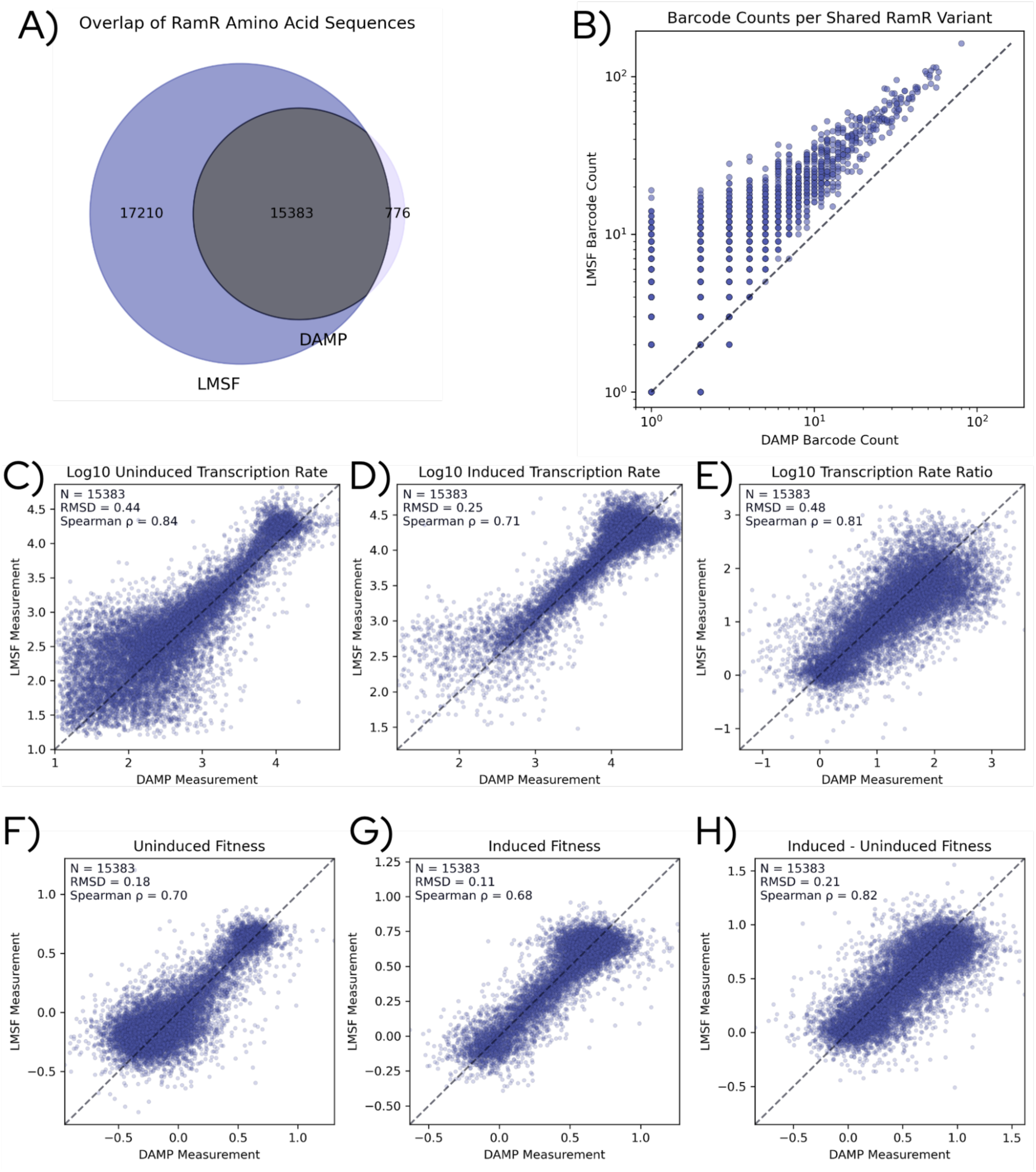
Corresponding main text Figure 2 for RamR, with comparison of calibrated (C–E) and uncalibrated (F–H) measurements across sites. Calibrated measurements show strong agreement, with Spearman correlations of 0.84, 0.71, and 0.81 and RMSD values of 0.44, 0.25, and 0.48 for uninduced, induced, and ratio measurements, respectively. Uncalibrated measurements exhibit similar rank-order agreement (Spearman = 0.70, 0.68, 0.82) and lower RMSD values (0.18, 0.11, 0.21) due to differences in scale. Because the axes differ between calibrated and uncalibrated representations, RMSD values are not directly comparable; however, calibration preserves strong concordance while enabling interpretable, quantitatively scaled measurements between facilities.

**Supplemental Figure 8:**
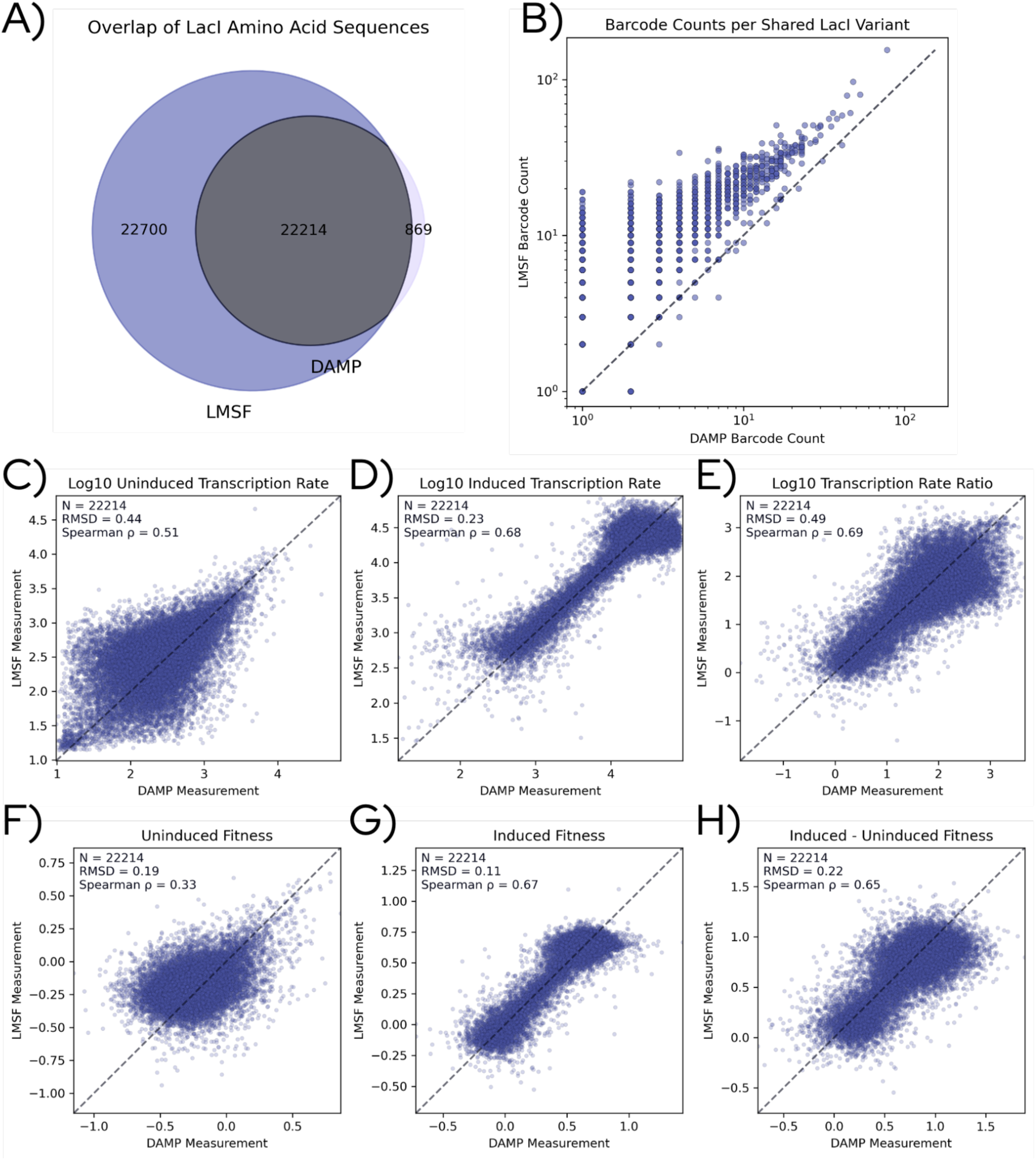
Corresponding main text Figure 2 for LacI, including uncalibrated measurements of fitness.

**Supplemental Figure 9:**
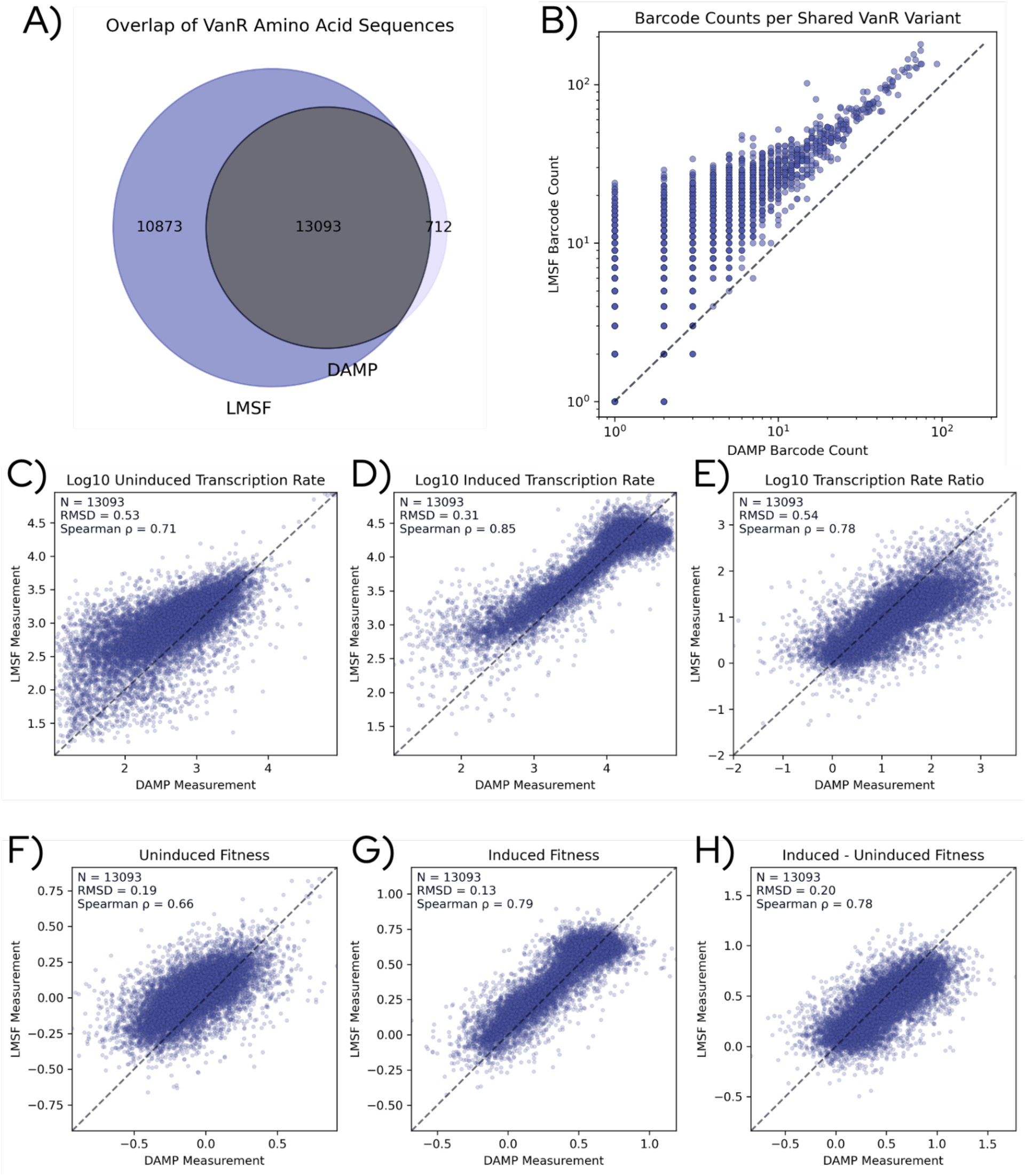
Corresponding main text Figure 2 for VanR, including uncalibrated measurements of fitness.

**Supplemental Figure 10:**
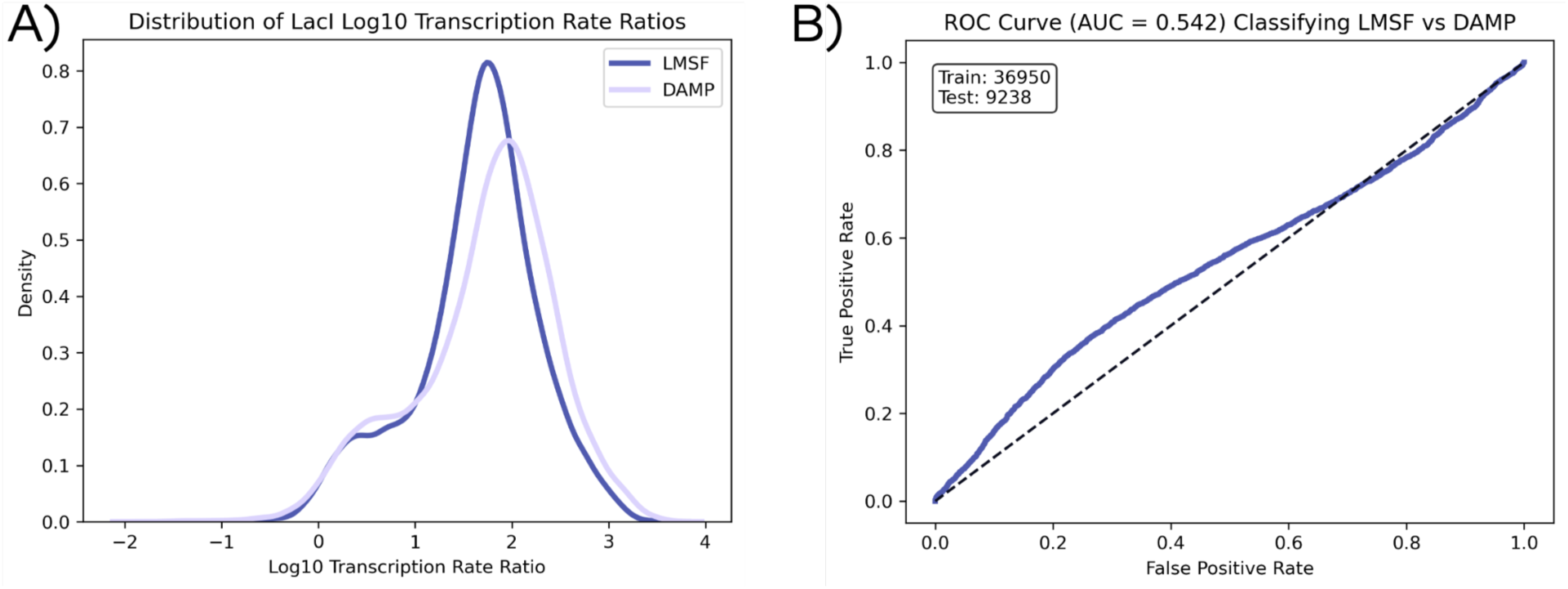
Corresponding main text Figure 3 for LacI.

**Supplemental Figure 11:**
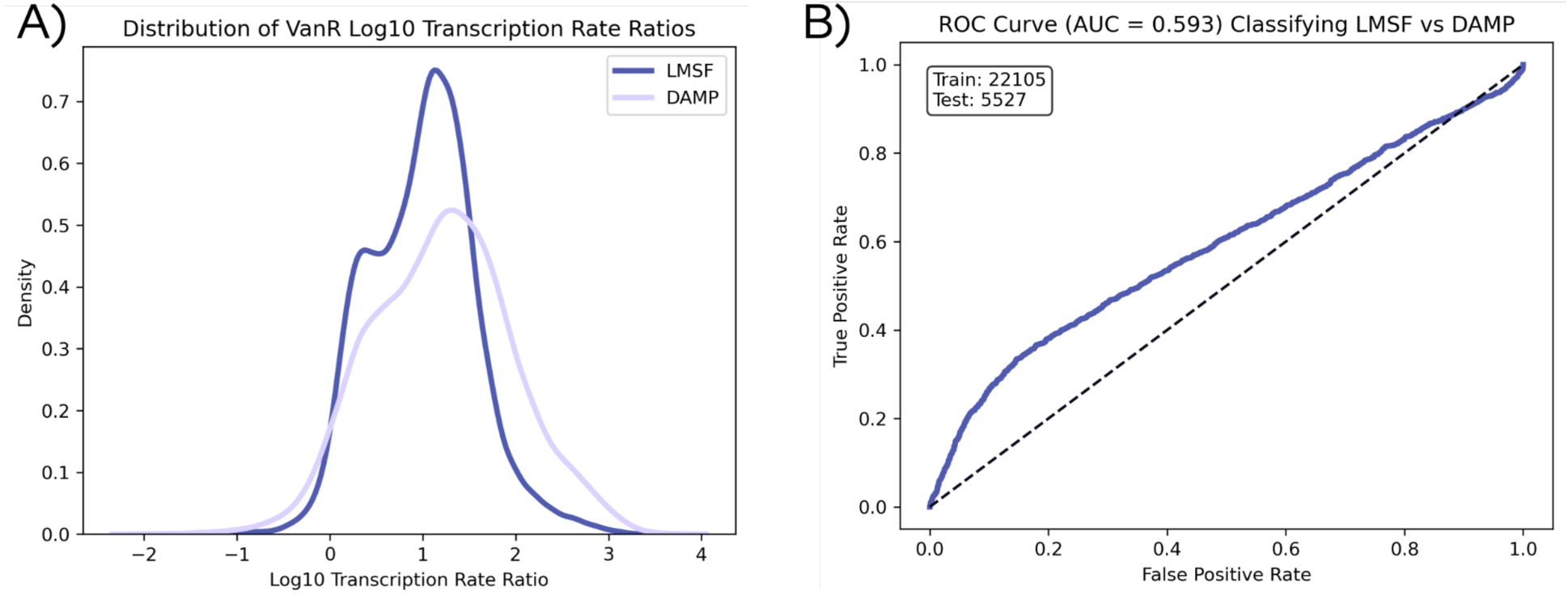
Corresponding main text Figure 3 for VanR.

**Supplemental Figure 12:**
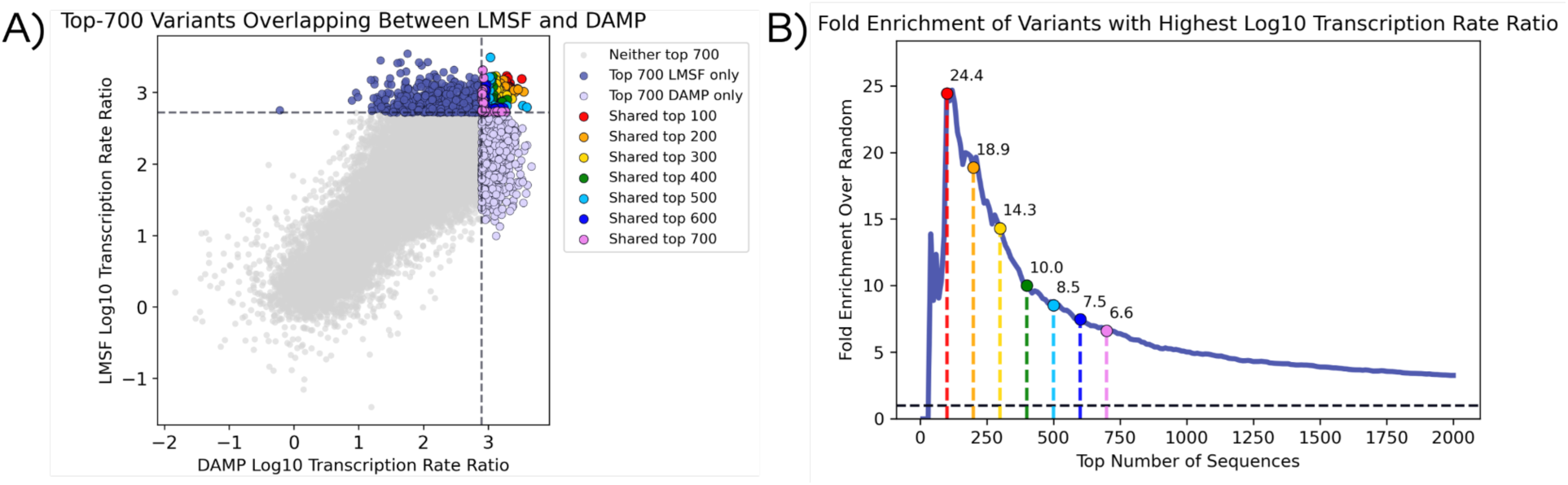
Corresponding main text Figure 4 for LacI.

**Supplemental Figure 13:**
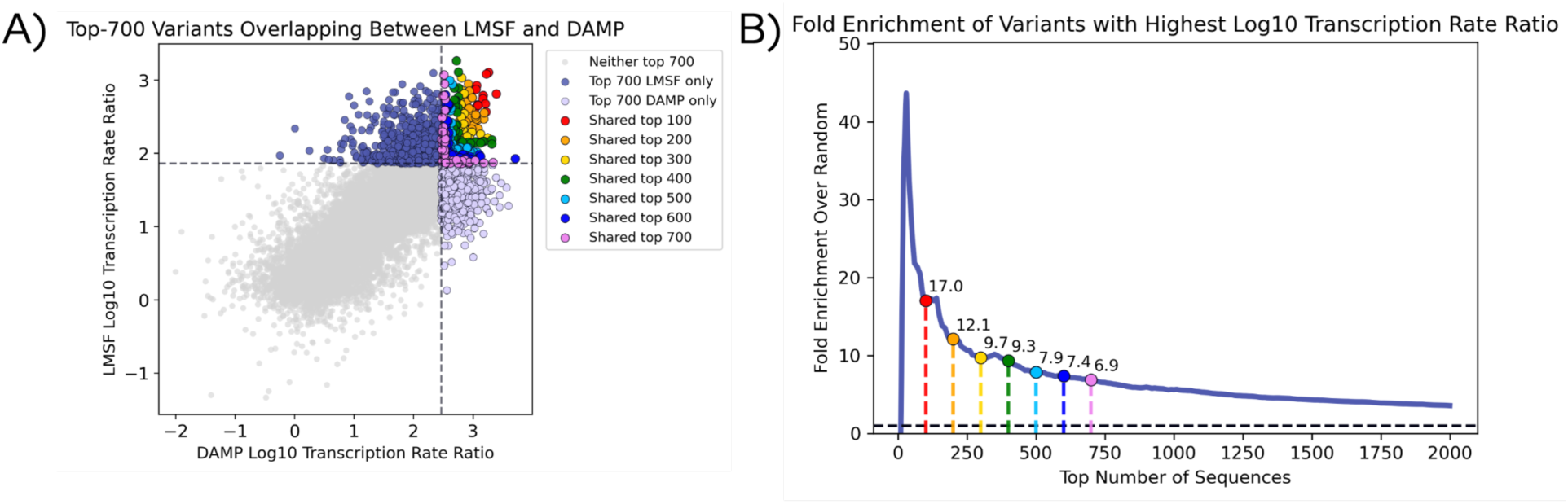
Corresponding main text Figure 4 for VanR.

**Supplemental Figure 14:**
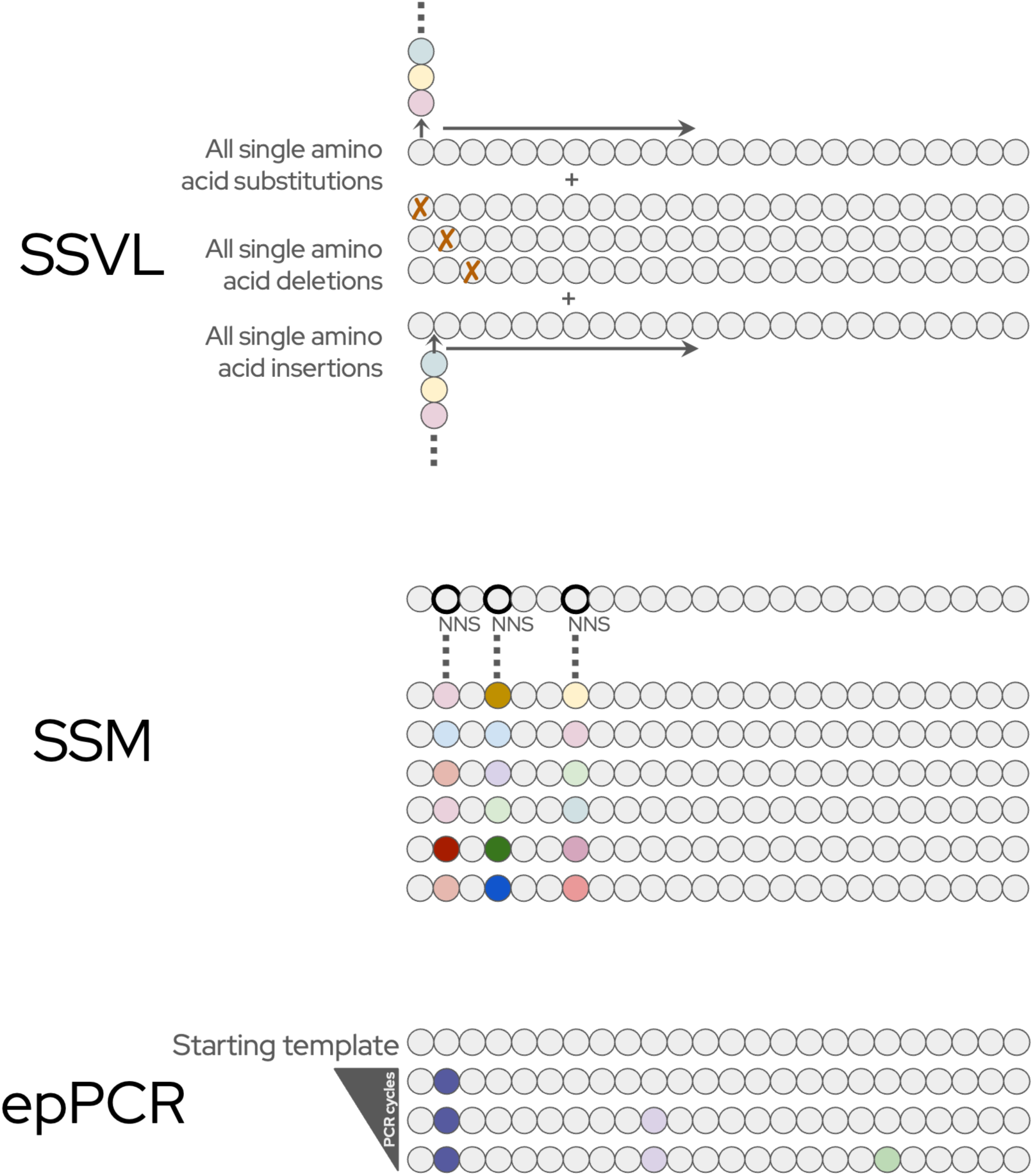
Library types used in this report. Here, we report data on site saturation variant (SSVL), site saturation mutagenesis (SSM), and error-prone PCR (epPCR) libraries. SSVLs include all single amino acid substitutions, deletions and insertions. SSVLs provide control over the final variant library but are limited to a single change per variant. SSM libraries introduce diversity via PCR with oligos containing degenerate NNS codons targeting 3 nearby amino acid residues. Each SSM library combinatorially randomizes 3 nearby residues. Three sets of NNS codons produces (20 x 20 x 20 =) 8000 unique protein variants and (32 x 32 x 32 =) 32,000 unique gene variants. SSM libraries allow targeted sites for random combinatorial mutations. Meanwhile, epPCR libraries are random but allow for many variants per library, spread out across the entirety of the gene. By assessing these 3 types of library for each transcription factor, we probe highly precise single mutations, targeted random combinatorial mutations of 3 proximal residues, and randomly generated multimutants.

